# Enzymatic DNA synthesis for digital information storage

**DOI:** 10.1101/348987

**Authors:** Henry H. Lee, Reza Kalhor, Naveen Goela, Jean Bolot, George M. Church

## Abstract

DNA is an emerging storage medium for digital data but its adoption is hampered by limitations of phosphoramidite chemistry, which was developed for single-base accuracy required for biological functionality. Here, we establish a *de novo* enzymatic DNA synthesis strategy designed from the bottom-up for information storage. We harness a template-independent DNA polymerase for controlled synthesis of sequences with user-defined information content. We demonstrate retrieval of 144-bits, including addressing, from perfectly synthesized DNA strands using batch-processed Illumina and real-time Oxford Nanopore sequencing. We then develop a codec for data retrieval from populations of diverse but imperfectly synthesized DNA strands, each with a ~30% error tolerance. With this codec, we experimentally validate a kilobyte-scale design which stores 1 bit per nucleotide. Simulations of the codec support reliable and robust storage of information for large-scale systems. This work paves the way for alternative synthesis and sequencing strategies to advance information storage in DNA.

## Main

DNA is a compelling data storage medium given its superior density, stability, energy-efficiency, and longevity compared to commonly used electronic media (*1*, *2*). Recent studies have demonstrated that digital data can be written in DNA, stored, and accurately read (*3*–*9*). However, the adoption of DNA for information storage remains limited due to its reliance on phosphoramidite chemistry, the method of choice for *de novo* DNA synthesis. This chemistry, designed for synthesizing DNA with single-base accuracy for biological applications, comprises several lengthy reactions (**Fig. S1A**) and costly reagents (*10*). Thus, alternative processes which could improve DNA synthesis cost and speed are highly desirable for large-scale information storage in DNA.

Here, we devise a *de novo* DNA synthesis strategy and a digital codec designed specifically for information storage. In contrast to DNA synthesized for biological functionality, DNA for information storage does not require single-base precision and accuracy. For synthesis, we harness a template-independent DNA polymerase, a protein evolved to rapidly catalyze the linkage of naturally occurring nucleotide triphosphates (dNTPs) under non-toxic biocompatible conditions. We encode information in transitions between non-identical nucleotides, rather than single nucleotides. We demonstrate that enzymatic synthesis and tailored computational tools provide robust information storage, as assessed using batch (Illumina) and real-time (Oxford Nanopore) sequencing. Moreover, our projections indicate that our enzymatic synthesis strategy is cheaper than phosphoramidite chemistry and may reduce reagent costs by orders of magnitude, facilitating the adoption of DNA as a storage medium.

## Enzymatic DNA synthesis

We chose to use terminal deoxynucleotidyl transferase (TdT), a template-independent DNA polymerase which rampantly and indiscriminately adds dNTPs to the 3’ termini of DNA strands (*11*–*15*). As such, TdT is largely used in reactions where one nucleotide triphosphate is added to indeterminate lengths (*16*–*18*). Inspired by previous work (*19*–*21*), we sought to leverage apyrase, which degrades nucleotide triphosphates into their TdT-inactive diphosphate and monophosphate precursors. By competing with TdT for nucleotide triphosphates, apyrase effectively limits DNA polymerization. We thus created and optimized a mixture containing a tuned ratio of these two enzymes such that a nucleotide triphosphate is added at least once to each strand by TdT before being degraded by apyrase (**Figs. 1, S2-S5, Supplementary Text 1.1-1.2**). We further determined the lowest nucleotide triphosphate concentrations required such that adding a series of nucleotides results in stepwise increases in the length of synthesized DNA (**Supplementary Text 1.3, Figs. S6-S7)**.

**Figure 1.**
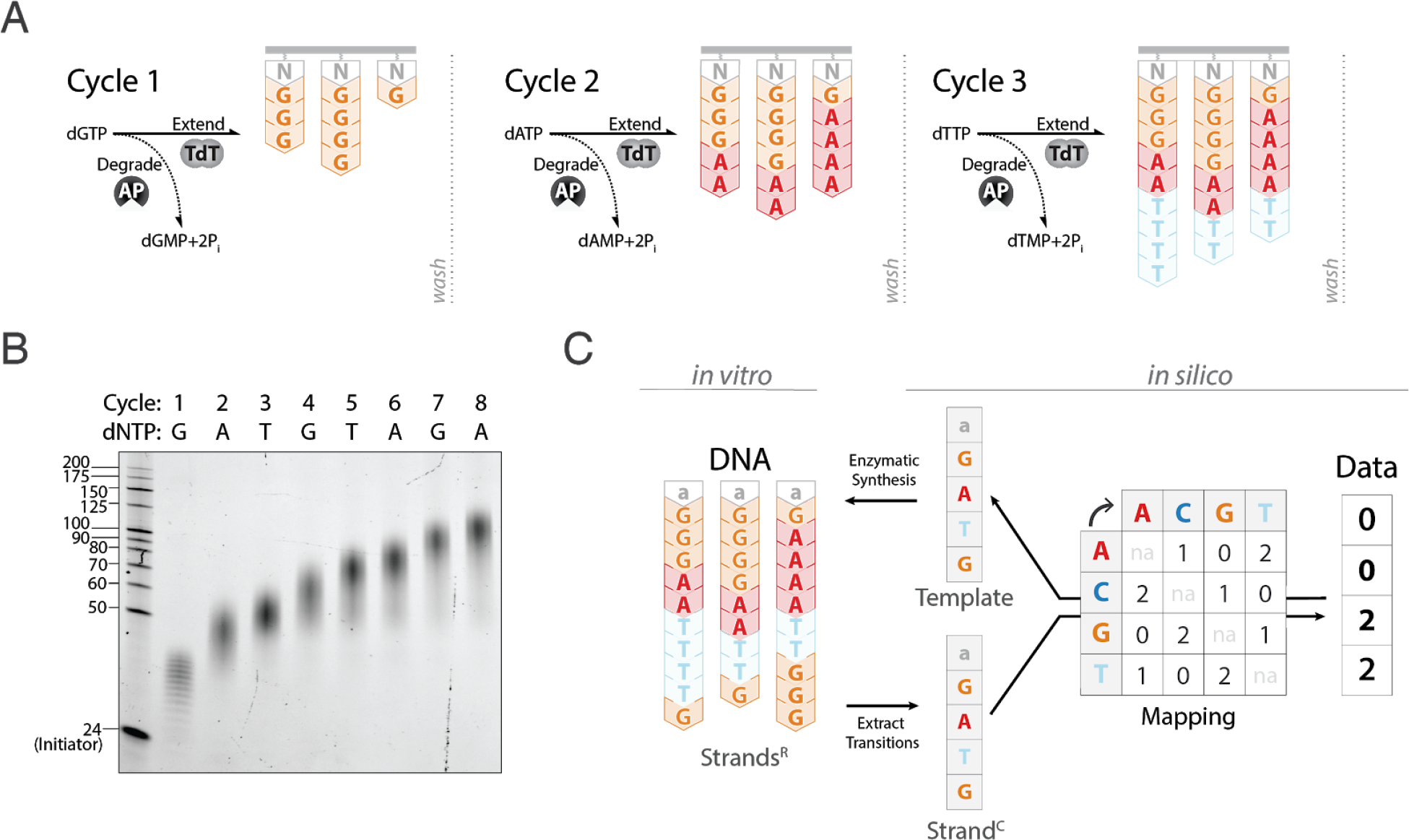
An enzymatic synthesis strategy for storing information in DNA. (A) Schematic depiction of a series of enzymatic synthesis reactions consisting of an oligonucleotide initiator (N, gray), terminal deoxynucleotidyl transferase (TdT) and apyrase (AP). The initiator is tethered to a solid support. In each cycle, TdT catalyzes the addition of a given nucleotide triphosphate to the 3’ end of all initiators while apyrase degrades the added substrate to limit net polymerization. A wash can be performed at the end of each cycle to remove reaction byproducts or to facilitate downstream processes. (B) DNA strands synthesized for each of eight consecutive synthesis cycle, as shown on 15% TBE-urea gel. The initiators were not tethered to a solid support and no wash was performed between cycles. The first lane is a single-stranded DNA size marker which includes 24 nucleotide long initiator oligonucleotide. (C) A schema for interconversion of DNA and information. Raw strands (strands^R^) represent enzymatically-synthesized DNA. A compressed strand (strand^C^) represents a sequence of transitions between non-identical nucleotides. Transitions between nucleotides, starting with the last nucleotide of the initiator (as an example N = ‘a’, gray) are mapped from the compressed strand to digital data in trits. If a strand^C^ is equivalent to the template sequence, all desired transitions are present and the information stored in DNA is retrieved.

Our enzymatic synthesis strategy requires few components to rapidly polymerize DNA (**Fig. 1A, S1B**). The core of the reaction is a mixture of TdT, apyrase, and short oligonucleotide initiators. Upon addition of a nucleotide triphosphate substrate, TdT extends the initiators until all added substrate is degraded by apyrase. We define the number of polymerized nucleotides as ‘extension length’. Subsequent nucleotide triphosphates are added to continue the synthesis process. While the extension length for each added nucleotide triphosphate may vary, the resulting population of synthesized strands all share the same number and sequence of nucleotide transitions (**Fig. 1B**).

We chose to encode information as transitions between non-identical nucleotides (**Fig. 1C**). Given three possible transitions for each nucleotide, we use trits rather than bits to maximize information capacity. To convert information to DNA, information in trits is mapped to a template sequence which represents the corresponding transitions between non-identical nucleotides starting with the last nucleotide of the initiator. Enzymatic DNA synthesis of each template sequence produces ‘raw strands’, or strands^R^, which can be physically stored. To retrieve information stored in DNA, strands^R^ are sequenced and transitions between non-identical nucleotides extracted, resulting in ‘compressed strands’, or strands^C^. If a strand^C^ is equivalent to the template sequence, the strand (compressed or raw) is considered ‘perfect’ and the information is retrieved by mapping the sequence of non-identical nucleotides back to trits.

To demonstrate storage of information, we encoded and synthesized “hello world!”, a message containing 96-bits of ASCII data (**Fig. 2A**). We split this message into twelve individual 8-bit characters, and prefixed each character’s bit representation with a 4-bit address to denote its order. These 144 total bits of information, including addressing, were also expressed in trits and mapped according to nucleotide transitions (**Fig. 1C**), resulting in twelve eight-nucleotide template sequences (**Table S2**). We synthesized all twelve template sequences (H01-H12) in parallel on bead-conjugated initiators while washing every two cycles. Following the last synthesis cycle, all strands^R^ were ligated to a universal adapter, PCR amplified, and stored as a single pool (**Methods**).

**Figure 2.**
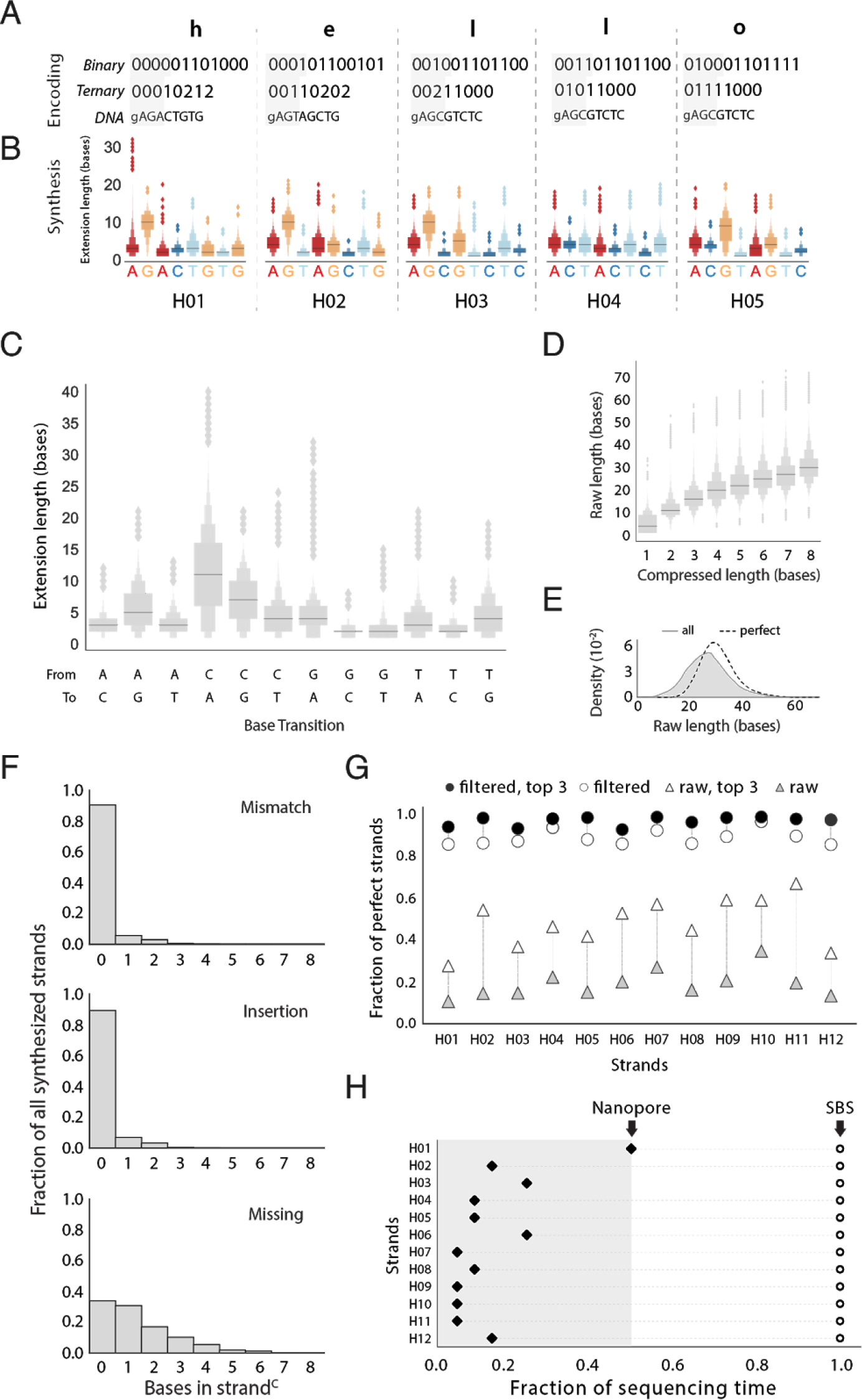
Demonstration of information storage in DNA using enzymatic synthesis. (A) The message “hello world!” was encoded in twelve template sequences, H01-H12, each representing one character. Transitions between nucleotides starts with the last base of the initiator, which is labeled ‘g’. A header index (shaded gray) denotes strand order. Only results from H01-H05 are shown (see **Fig. S9**). To encode each character, its respective ASCII decimal value, prefixed with an address is represented in base 2 (binary) or in base 3 (ternary) (see **Table S2**), mapped to transitions (see **Fig. 1C**), resulting in template sequences with nucleotides to be synthesized (capitalized). (B) Extension lengths for each base from (A). Only perfect strands^R^, those whose strand^C^ is equivalent to a template sequence, are considered here. Synthesis was performed with initiators tethered to beads and sequencing performed on the Illumina platform. (C) Distribution of extension lengths for each nucleotide transition, combined across all positions from all perfect strands. (D) Stepwise increases in strand^R^ length with an increasing strand^C^ length for all synthesized strands of H01-H12. (E) Distribution of all strand^R^ lengths. Distributions are derived via kernel density estimation for all synthesized strands (‘all’, gray shading) and a subpopulation of strands that contain all desired transitions (‘perfect’, dotted line). (F) Bulk error analysis for all synthesized strands of H01-H12. All strands^C^ were aligned, by Needleman-Wunsch, to their respective template sequences, and the number of mismatches, insertions, and missing nucleotides were tabulated. (G) Information retrieval with *in silico* filtering. Fraction of perfect strands^C^ are shown before (triangles) or after filtering (circles). Fraction of perfect strands^C^ is shown for all sequences (white) or only the top 3 most abundant sequences (black). (H) Information retrieval by different sequencing platforms. Streaming nanopore sequencing (Oxford, filled diamonds) was compared to batch sequencing-by-synthesis (Illumina, open circles). Each dot indicates the fraction of sequencing run at which each strand is robustly retrieved (100% correct with 99.99% probability). Arrows denote the fraction of the sequencing run at which all data is robustly retrieved using each platform.

## Data retrieval and error analysis

We used Illumina sequencing to read out our synthesized strands^R^ and to assess the information stored in corresponding strands^C^(**Methods**). We started by analyzing the perfect strands. We found that the extension length for each nucleotide varied based on the type of transition (**Fig. 2B, S9, Table S3**). As a result, perfectly synthesized strands for each template sequence may be of variable raw length. Additionally, when extension lengths were compiled for each nucleotide across strands and positions based on type of transition, we observed that these lengths were qualitatively consistent between bead-conjugated (**Fig. 2C**) and freely-diffusing initiators (**Fig. S6, Supplementary 1.3**). For example, the median extension lengths of C when following A, T, or G were among the lowest. Conversely, the median extension lengths for A, T, and G when following C were among the highest. Considering all synthesized strands, we found stepwise increases in the median raw lengths with an increasing number of non-identical nucleotides (compressed strand length), indicating controlled polymerization for the population of strands over multiple cycles (**Fig. 2D**). However, compared to a median length of 30 nucleotides for all perfect strands^R^, the median length for all synthesized strands^R^ was 26 bases, suggesting that not every strand polymerized the added nucleotide triphosphate in each cycle (**Figs. 2E, S10**).

To identify the types and magnitude of synthesis errors in this system, we aligned all synthesized strands^C^ to their respective template sequences and tabulated the number of missing, mismatched, and inserted nucleotides (**Figs. 2F, S11, Methods**). While multiple alignments exist for several imperfect strands^C^, which ambiguate the exact position of errors, the type of error for each strand^C^ can be distinguished. Our analysis indicates that 9.5% of strands^C^ contained one or more mismatches, 10.7% contained one or more insertions, and 66.1% contained one or more missing nucleotides. Thus, the dominant type of error is missing nucleotides in a strand^C^, which corresponds to a strand that did not get extended by an added nucleotide triphosphate in at least one synthesis cycle.

In spite of synthesis errors, we retrieved information from our pool of synthesized DNA strands^C^ by applying a simple two-step *in silico* filter. As each template sequence is designed with a specific architecture (**Methods**), we first filtered synthesized strands^C^ by length and presence of a terminal ‘C’. Using this filter, the fraction of perfect strands for all template sequences (H01-H12) increased from an average of ~19% to an average of ~89% (**Fig. 2G**). We then selected for the most abundantly synthesized strand^C^ variant in this subset to retrieve data.

Finally, we show that quick access to information stored in DNA may be accomplished with real-time sequencing. While the Illumina platform sequences all DNA strands in parallel and reports the outcome in batch, the Oxford Nanopore platform offers asynchronous sequencing by translocation of DNA strands through independent nanopores, and streams the outcome. As a result, sequencing can be terminated as soon as data is retrieved and remaining reagents provisioned for later use.

To demonstrate the advantage of real-time information retrieval, we first sequenced DNA strands^R^ synthesized for H01-H12 using an entire MinION flowcell (Oxford Nanopore) and observed that the most abundant species, an average of 49.9% of filtered strands^C^, were perfectly synthesized (**Fig. S12A**). This is largely consistent with results from Illumina sequencing, with the slight decrease likely due to errors currently inherent to state-of-the-art nanopore sequencing (*22*). With these experimental results, we then performed simulations to determine the fraction of sequencing resources required for robust data retrieval from each of the twelve template sequences H01-H12 with at least 99.9% probability. We simulated repeated trials which, at a given fraction of the total sequencing run, randomized the translocation time of each DNA strand^R^ through the nanopore and assessed whether data could be retrieved (**Methods**). Our simulations indicate that only half of the total sequencing resources were needed to robustly retrieve data from DNA using Oxford Nanopore compared to Illumina (**Figs. 2H, S12B, Table S5**).

More broadly, nanopore sequencing can enable faster and more efficient information retrieval from strands synthesized with our enzymatic strategy. Currently, DNA translocation rates are slowed through nanopores for accurate single-base sequencing. This rate may be increased since it is, in principle, easier to detect transitions between non-identical nucleotides, each with extension lengths greater than one (*23*–*26*). Furthermore, through selective sequencing (*27*), nanopores could reject strands corresponding to already recovered sequences in favor of strands for remaining template sequences. Such an approach may be accomplished by detection of each strand’s address. These alternative design parameters may inspire the development of sequencing technologies that are faster, more affordable, and specifically designed for DNA information storage.

## Coded strand architecture

We have established that data can be stored in enzymatically-synthesized DNA and retrieved by *in silico* filtering for perfectly synthesized DNA strands. However, perfect strands^C^ may not be required for data retrieval. Imperfectly synthesized strands^C^, which carry partial information, may be used to reconstruct template sequences if nucleotide errors occur in different locations. We thus sought to develop a codec for robust data retrieval which leverages the diversity of imperfectly synthesized strands^C^ for template sequence reconstruction. The core of our codec relies on three elements: (i) A coded strand architecture which includes synchronization nucleotides to facilitate error localization, (ii) Sufficiently diverse strands^C^ produced by synthesis, and (iii) Sequence reconstruction from strands^C^ with statistical inference based on mathematical models of synthesis. Our codec models information storage in DNA as a communications channel to enable correction of errors accumulated from synthesis, storage, and sequencing (**Fig. 3A**).

A key feature of this codec is the addition of synchronization nucleotides which are interspersed between information-encoding nucleotides (**Fig. 3B**). These nucleotides act as a scaffold that aids the reconstruction of a template sequence from DNA strands^C^ that may contain errors as a result of missing, mismatched, and inserted nucleotides. As an example, consider a template sequence of 8 nucleotides (CTCGTGCT) and two synthesized DNA strands^C^(CTCTGC and TCGTCT), each with two missing nucleotides. Without a scaffold, data cannot be retrieved since three equally valid reconstructions are possible. In contrast, a scaffold constrains the number of possible sequences to one, allowing data retrieval from otherwise unusable DNA strands^C^. Accordingly, our codec includes a module for encoding information in template sequences which incorporates synchronization nucleotides (**Supplementary Text 2.5**).

**Figure 3.**
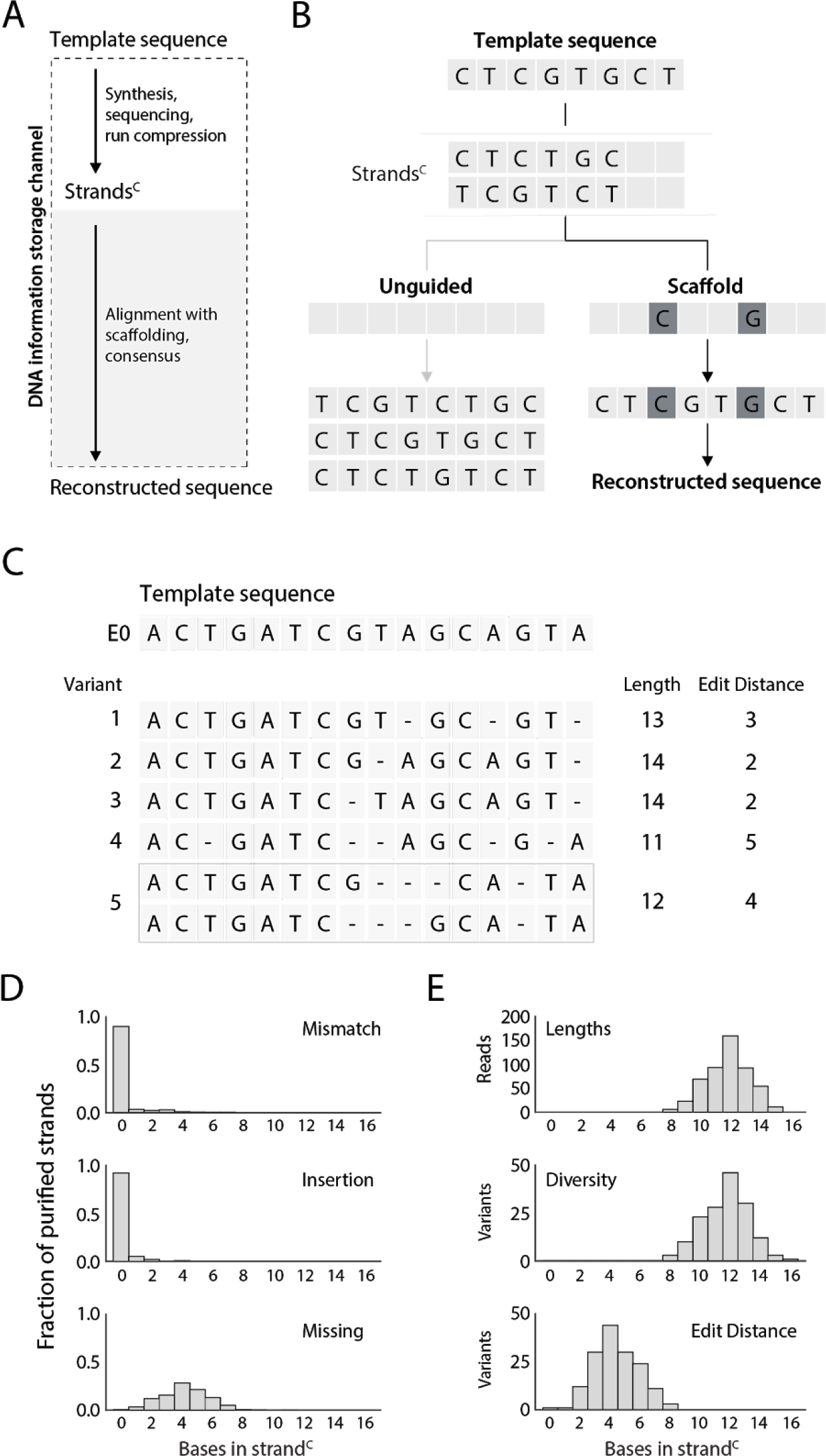
Coded strand architecture for sequence reconstruction. (A) A DNA information storage channel. Data is converted to template sequences, synthesized (yielding strands^R^), and can be stored *in vitro*. Retrieval starts with sequencing, then transitions of non-identical nucleotides are extracted *in silico* to form strands^C^. Data retrieval occurs when the template sequence and reconstructed sequence are equivalent. Errors which occur in the synthesis and sequencing steps can be modeled as a communications channel. (B) A coded strand architecture, ‘scaffold’, enables data retrieval from strands^C^ that are missing nucleotides, whereas an ‘unguided’ reconstruction results in multiple possible solutions. Synchronization nucleotides (dark gray boxes) localize errors to yield a single reconstructed sequence. (C) A 16-base transition sequence, E0, is synthesized and sequenced with Illumina. Examples of diverse strands^C^ produced by synthesis of E0. Strands^C^ are aligned, by Needleman-Wunsch, to the template. Ambiguous alignments can exist depending on the location and number of missing nucleotides within a strand^C^. (D) Error analysis for purified strands of E0. Synthesized strands were purified *in silico*, by filtering for strands^R^ between 32-48 bases in length, and corresponding strands^C^ were aligned by Needleman-Wunsch to the E0 template. For each alignment, the number of mismatches, insertions, and missing nucleotides were tabulated. (E) Evaluating the diversity of synthesized strands. The number of sequencing reads for each length of strand^C^ was tabulated. Diversity was evaluated as the number of unique variants at each length of strand^C^ and the Levenshtein edit distance was computed with respect to the E0 template. The set of 802 purified strands contains 2 perfect strands.

To reconstruct missing nucleotides from strands^C^ by scaffolding, the population of synthesized DNA strands for a desired sequence must be sufficiently diverse. That is, if the same nucleotide is missing systematically across all strands, then it cannot be retrieved without additional forms of error correction. To analyze the diversity generated from our enzymatic process, we synthesized a longer 16-nucleotide template sequence (called E0), which contains 12 unique transitions between nucleotides to mitigate ambiguous alignments (**Fig. 3C**). We performed *in silico* size selection of strands^R^ ranging 32 to 48 bases in length, assuming that the 16 template nucleotides were synthesized with an average extension length of two to three bases (**Fig. S13A**). We analyzed this purified set by aligning the corresponding strands^C^ to the E0 template and observed that missing nucleotides were the predominant form of error, in line with our previous analyses (**Fig. 2F**), but occurred in different positions (**Figs. 3C,D, S13B**).

We also assessed diversity by analyzing the lengths, number of variants, and Levenshtein edit distances of strands^C^ from our purified set (**Fig. 3D, S13C**). We observed that the median strand^C^ length was 12 nucleotides and the maximal number of variants occurred at this length. We also calculated the Levenshtein edit distance (*28*), which summarizes the number of single-nucleotide edits required to repair a strand^C^ to the desired E0 sequence. The median edit distance for these variants was four, indicating that synchronization nucleotides could be placed approximately every three or four nucleotides to recall missing strand^C^ nucleotides from diversely synthesized strands (**Supplementary Text 3**). Together, these analyses provide the data for creating the statistical inference element of our codec.

We next set out to establish the statistical inference and mathematical models that would enable the reconstruction of a template sequence from a population of diverse but imperfect strands^C^. For efficient reconstruction, we adapted a statistical framework known as *maximum a posteriori* (MAP) estimation (*29*). To utilize this framework, we built a Markov model to describe the synthesis of a strand^C^ with error probabilities for mismatches, insertions, and missing nucleotides, derived from our analyses of the purified set of E0 strands^C^(**Fig. S16A**). These state probabilities can be used to score all possible reconstruction solutions consistent with a scaffold, considering mismatches and insertions in addition to missing nucleotides (**Fig. S17, Supplementary Text 2.7**). Our calculations provide a probability of occurrence for each nucleotide at each position. Ultimately, a consensus can be obtained to indicate the most probable nucleotide per position (**Supplementary Text 2.7**), which ideally yields a reconstructed sequence that is equivalent to the template sequence.

Finally, we sought to experimentally verify our codec design by encoding and synthesizing the message “Eureka!” as four template sequences, E1-E4 (**Fig. 4A, Supplementary Text 2.5**). Each template sequence contained a 2-bit address to delineate its order, and 14 bits of data. These 16 bits are encoded in a template sequence of 16 nucleotides, which includes four synchronization nucleotides, resulting in 1 bit stored per nucleotide (**Supplementary Text 2.5, Fig. S15B**). Sequences E1-E4 carry a total of 64 bits of information including addressing, and were synthesized in parallel on beads with a wash every cycle. Following the last synthesis cycle, strands were ligated to a universal adapter, PCR amplified, and stored as a single pool (**Methods**).

**Figure 4.**
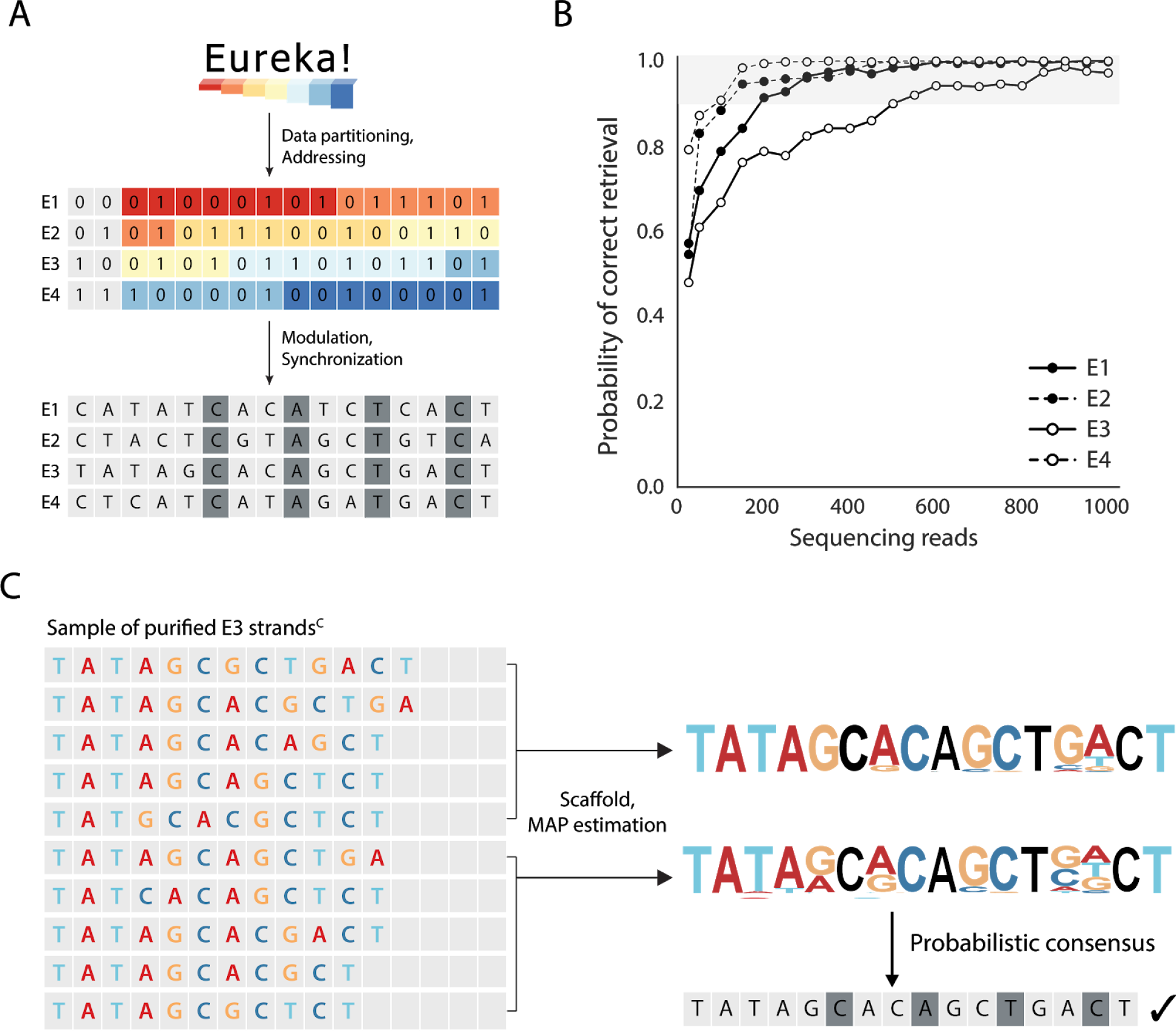
Coded strand architecture for robust information storage. (A) The message “Eureka!” was encoded and partitioned into four template sequences, E1-E4. Each sequence stores a 2-bit address and 14 bits of data. These bits are mapped to a template sequence of 16 nucleotides, which includes four synchronization nucleotides (dark gray). Synthesis was performed with initiators tethered to beads and sequencing performed on the Illumina platform. (B) Retrieving information from E1-E4. Synthesized strands^R^ were sequenced using the Illumina sequencing-by-synthesis (SBS) platform and purified *in silico* based on raw length of 32-48 nucleotides (**Methods**). The decoding accuracy for each sequence is defined as the probability of 100% correct data retrieval for a given number of reads, estimated over 500 decoding trials. Each trial is based on a randomly drawn set of purified strand^C^ variants. A 90% decoding accuracy (gray band) is considered sufficient for robust data retrieval, and this accuracy could be further reinforced by other codec modules. (C) Decoding of E3. A set of 10 DNA strands^C^ is decoded as two sets of five strands^C^. The decoder uses MAP estimation and a scaffold to determine the probability for each of the four nucleotides at every position. The decoded sequence is a probabilistic consensus of the reconstructed sequences from MAP estimation and successfully retrieves the data stored in E3.

We reconstructed our stored message “*Eureka!*” by using only implicit error correction provided by the diversity generated from enzymatic synthesis. We performed *in silico* size selection of all strands^R^ of length 32-48 nucleotides (**Fig. S18**). This set of 4521 purified strands^R^ contained 31 perfect strands (**Fig. S20B**). We then attempted to reconstruct template sequences by MAP estimation with scaffolding and probabilistic consensus (**Supplementary Text 2.7**). We found that 10 strand^C^ variants, each with an error tolerance of ~30% as a result of missing an average of 4 or 5 out of 16 nucleotides, could accurately reconstruct a template sequence (**Fig. S26**). We then assessed the number of sequencing reads required for a 90% probability of data retrieval. We found that all four template sequences were robustly reconstructed with 200, 150, 500, and 100 sequencing reads for E1-E4 respectively, with a median of 175 reads (**Fig. 4B**). Sequence E3 required the most sequencing reads for reconstruction as synthesized strands contained one extra edit on average in comparison to synthesized strands for other template sequences (**Figs. S19-S20)**. We also found that MAP estimation was a more robust decoding algorithm than our previous two-step filter for H01-H12, requiring fewer reads for data retrieval (**Fig. S26**). These results show that our codec can accurately reconstruct data without requiring perfectly synthesized DNA strands.

## Scalable Codec for DNA Information Storage

Our experimental results demonstrate that byte- and kilobyte-scale storage systems can be achieved if a sufficient number of strands are synthesized (**Fig. 5A**). Specifically, our “hello world!” experiment stored 12 bits per template sequence. This is sufficient for a 256-byte maximum storage system where 11 bits are used for addressing 2,048 total template sequences, each with 1 bit of data. In contrast, our “Eureka!” experiment stored 16 bits per template sequence. This allows for a 4-kilobyte maximum storage system, where 15 bits are used for addressing 32,768 total template sequences, each with 1 bit of data (**Table S7, Supplementary Text 2.2**).

**Figure 5.**
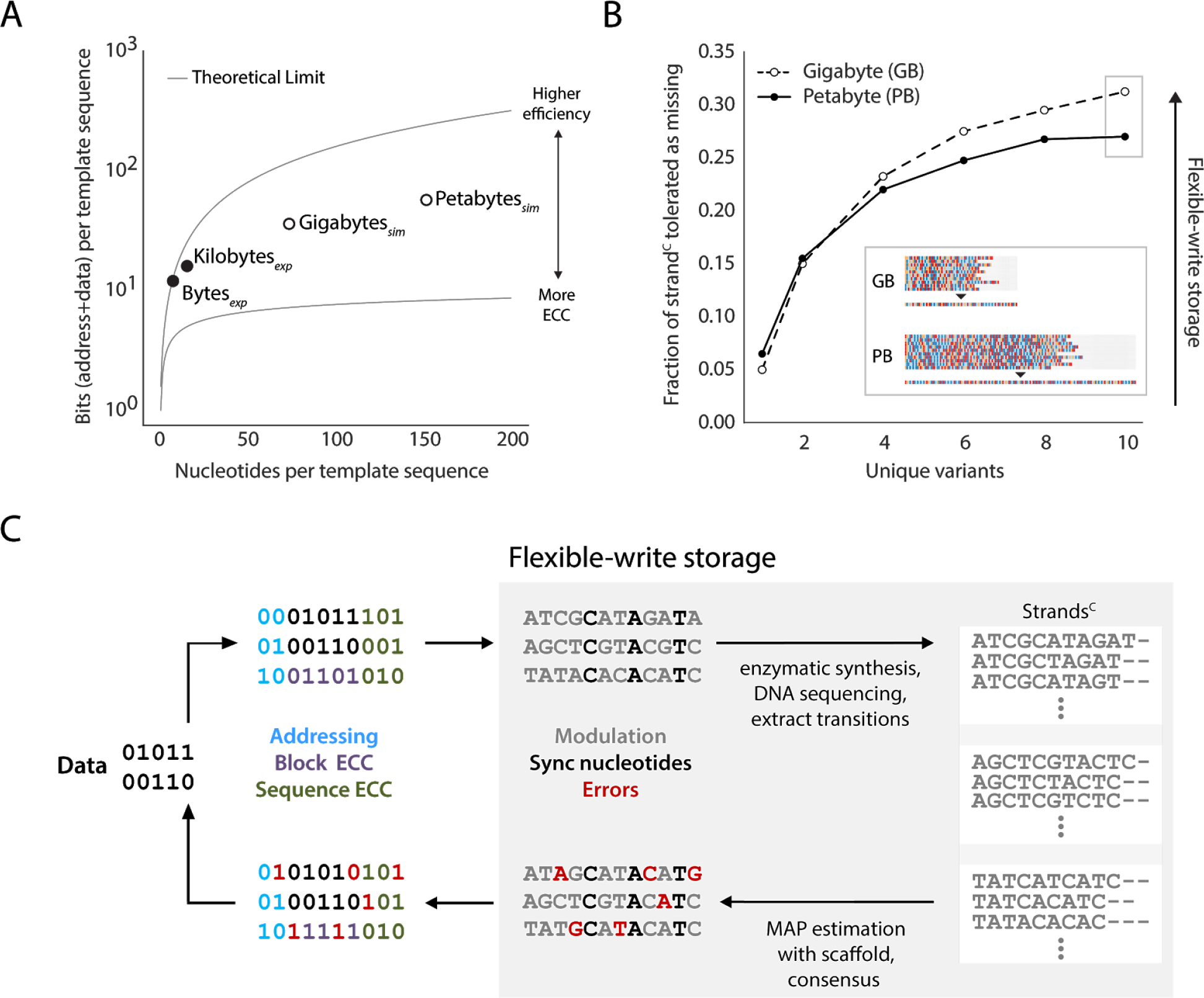
A roadmap for scaling DNA storage systems. (A) Efficiency of storage for experimental and simulated systems. Experimental systems (black) include storing 12 bits in 8-nucleotide template sequences, and 16 bits in 16-nucleotide template sequences. Simulated maximum storage systems (white circles) include gigabyte scale which stores 36-bits in a 74-nucleotide template sequence, and petabyte scale which stores 57-bits in a 152-nucleotide template sequence. The amount of bits stored per sequence is dependent on the amount of error-correction codes (ECC) that are applied. Reducing ECCs increases the efficiency rate of storage. The upper bound theoretical limit represents a maximum efficiency of storage of ~1.58 bits per transition between non-identical nucleotides (**Supplementary Text 2.5.3**). The lower bound theoretical limit represents the minimum number of bits per template sequence that must be stored for only addressing (**Supplementary Text 4.2**) (B) Flexible-write storage is enabled by a codec which harnesses diversely synthesized strands. The decoding pipeline supports robust data retrieval from synthesized strands with a significant percentage of errors. Inset: with ten strand^C^ variants, each with ~30% missing nucleotides, the correct decoded sequence can be reconstructed for both gigabyte- and petabyte-scale maximum storage capacities. (C) A system architecture for storing information in enzymatically-synthesized DNA. A bitstream is partitioned into rows, each augmented with an address to delineate its order for reassembly. An ECC such as a Bose–Chaudhuri–Hocquenghem (BCH) code can be applied to each row, or an ECC such as a Reed-Solomon (RS) code can be applied across multiple rows, to protect data from errors (**Supplementary Text 2.4**). Modulation consists of mapping sequences of bits to template sequences, which includes synchronization nucleotides. Enzymatic synthesis then produces multiple diverse strands^C^ per template sequence. The resulting strands^C^ are used for sequence reconstruction based on MAP estimation and probabilistic consensus. Subsequently, the reconstructed sequence is demodulated into bits. Error-correction is applied to ensure data retrieval.

We next assessed the scalability of our DNA storage codec for gigabyte- and petabyte-scale storage through simulation, under the assumption that the requisite number of DNA strands for each could be produced (**Supplementary Text 4**). Increased storage capacity requires more nucleotides per template sequence for additional address space, synchronization nucleotides, and data. Specifically, we simulated storing 36 bits, including data and address, in a 74-nucleotide template sequence and similarly, 57 bits in a 152-nucleotide template sequence to obtain gigabyte- and petabyte-scale systems, respectively (**Fig. 5A**). For these simulations, randomly generated data were partitioned, mapped to template DNA sequences, and synthesized strands^C^ were generated *in silico* using our Markov model for a wide range of synthesis accuracies (**Methods**). We provided an additional error-correction code (ECC) to each template sequence to ensure accurate reconstruction (**Supplementary Text 2.4**). We found that data could be accurately retrieved from our simulated synthesized strands with our decoding pipeline (**Fig. S28, Supplementary Text 2**). Our efficiency rates, calculated as bits stored per template nucleotide, may be considered competitive to those demonstrated in prior systems, considering that the bottleneck for attaining large-scale storage capacities is the massive parallelization of affordable synthesis reactions (**Supplementary Text 4**). With improvements to synthesis accuracy, our efficiency rates can be increased towards the theoretical maximum of ~1.58 bits per transition between non-identical nucleotides (**Fig. S27, Supplementary Text 2.6**).

To assess the robustness of our digital codec at higher storage capacities, we performed repeated decoding trials with many sets of strands^C^ synthesized *in silico* and measured the probability of data retrieval (**Methods**). Our simulations indicate that if at least 10 unique strand^C^ variants per sequence are available, then each variant could tolerate on average ~30% missing strand^C^ nucleotides (**Fig. 5B**). We found qualitatively similar results when simulated strands^C^ also included mismatch and insertion rates exceeding those observed experimentally, further illustrating the robustness of our codec (**Figs. S28**). Overall, our codec is able to resolve several types of errors, including missing nucleotides in synthesized strands^C^, which would otherwise drastically reduce information storage capacities (*30*, *31*). These simulated results affirm our experimental finding that data retrieval does not require perfectly synthesized strands.

Our scalable codec architecture consists of encoding and decoding frameworks to extract information from diversely synthesized DNA strands (**Figs. 5C, S30**). The encoder consists of several core components: (i) Partitioning of data into ordered rows of bits; (ii) Prefixing of rows with addresses; (iii) Error correction per row of bits via an error-correction code (ECC) per template sequence (e.g., Bose-Chaudhuri-Hocquenghem code), and error correction per block of rows via a block ECC (e.g., Reed-Solomon or Fountain code, (*5*–*7*)); (iv) Modulation to map rows of bits to template DNA sequences. All template sequences are subsequently synthesized enzymatically, resulting in a population of diverse DNA strands. Strands^R^ are read out by sequencing and corresponding strands^C^ are input to a decoder. The crucial first step of the decoding pipeline is MAP estimation aided by scaffolding, followed by probabilistic consensus. Multiple subsets of strands^C^ can be used for sequence reconstruction. Each reconstructed sequence need not be identical to the template sequence. After demodulation of the reconstructed sequence, the resulting bit sequence can be corrected by bit-level ECCs in the decoding pipeline to reinforce error-free data retrieval. Overall, our design harnesses the diversity of enzymatically-synthesized DNA strands and supports a flexible-write approach to provide a functional and robust storage system.

## Continued Improvements and Future Outlook

Taken together, our results show that information can be stored accurately in imperfectly synthesized DNA strands. However, it is important to point out current limitations of our approach in light of potential improvements and design optimizations. Assuming only size selected strands are stored for our kilobyte-scale design, our implementation incurs a 6-fold loss in volumetric density of information. This reduction is due to two factors: an extension length up to three bases per transition (~3-fold loss, **Fig. S18**) and an efficiency rate of storage of 1 bit per template nucleotide (~2-fold loss, **Supplementary Text 2.6**). This density loss is mild considering DNA’s thousandfold advantage over the projected fundamental density limit of flash drives (*2*, *3*, *6*) and may be addressable. The efficiency rate of storage may be increased as synthesis accuracy improves. Improved accuracy will also enable provisioning of TdT’s ability to add ~500 (**Fig. S4B**) to thousands (*13*) of nucleotides per strand^R^ to increase the number of strand^C^ nucleotides for increased storage capacities. On the other hand, extension lengths per template nucleotide may be considered a design optimization and tuned according to application demands, trading density for read-out speed and cost by specialized nanopore sequencing (*23*–*26*, *32*).

Currently, DNA for information storage is synthesized in a high-density array format with proprietary machines (*3*–*7*). We thus started to translate our bead-based process to a 2D array-based platform (**Fig. S31**). Using this prototype, we could produce perfectly synthesized strands for three 13-nucleotide template sequences (**Supplementary Text 5**). Analyses of the synthesized strands indicate similar error and diversity profiles to those observed using our bead-based process, indicating that our codec could be used to store information in DNA synthesized with this platform (**Figs. S32-S34**). Additional process engineering will be required to improve synthesis accuracy. For example, more stringent washing per cycle may reduce carryover of nucleotide triphosphates from previous cycles to further diminish the rate of substituted strand^C^ nucleotides. Optimization of reaction conditions to improve mixing or the use of more processive, rather than distributive, TdT mutants may reduce the rate of missing strand^C^ nucleotides (*33*). Additional efforts are also needed to improve automation and parallelization to increase DNA production for large-scale data storage.

Our enzymatic DNA synthesis strategy projects favorably in speed and cost relative to phosphoramidite chemistry. Specifically, assuming implementation on the same microarray instrument, we compared reagent costs for both processes as a function of feature size (reagent volume) (**Fig. S35, Table S6, Supplementary Text 6.2**). Our analyses indicate that our enzymatic synthesis strategy could already be cheaper as a drop-in replacement to phosphoramidite chemistry when using existing automation which synthesizes DNA strands in 15-30-micron features (**Fig. S35, Supplementary Text 6.2**). Further miniaturization, together with reductions to enzyme cost through recycling, provide a potential roadmap for overall reduction in reagent costs by several orders of magnitude (**Fig. S35**). In addition, the higher rate of enzymatic catalysis over chemical coupling and a lack of blocking moieties may shorten our synthesis cycle times compared to phosphoramidite chemistry, improving write speed and equipment amortization time (**Table S6, Supplementary Text 6.3**).

## Conclusion

In summary, we have presented an enzymatic synthesis strategy and tailored coding architecture for robust information storage in DNA. This storage solution is an alternative to prior studies which utilized phosphoramidite chemistry to produce DNA for information storage (*3*–*9*). Our approach offers potentially dramatic benefits to the cost and speed of synthesis and sequencing without requiring single-base accuracy. Additionally, our approach may alleviate biosecurity concerns associated with widespread DNA synthesis of genetic information, as genes are unlikely to be produced with this strategy. While this work illustrates DNA information storage *in vitro*, it could provide a foundation for development of *de novo* molecular recording systems *in vivo* (*34*–*37*). Further technological achievements, through industrial-grade automation, refined enzymatic reactions, and advanced coding systems, will improve the efficiency and accessibility of this platform and inform the design of a complete read and write system to advance large-scale DNA information storage.

## Acknowledgements

The authors would like to acknowledge John Aach, Nili Ostrov, and Javier Fernández Juárez for helpful discussions and comments on the manuscript, Calixto Saenz and the HMS Microfluidics/Microfabrication Facility, HMS Biopolymers Facility, and Dmitry Rodionov of Formulatrix, Inc. for technical support. This work was supported by funding from National Institutes of Health Grant R01-MH103910-02 (to GMC), Department of Energy Grant DE-FG02-02ER63445 (to GMC), and AWS Cloud Credits for Research program (to HHL, GMC).

## Author Contributions

HHL, RK, NG, JB, and GMC conceived the study. HHL, RK, and GMC developed enzymatic synthesis.NG developed the codec. HHL, RK and NG performed experiments and analyzed the data. HHL, NG, and RK wrote the manuscript. All authors reviewed and edited the manuscript. GMC supervised the study.

## Competing financial interests

All authors have filed patents related to this work.

